# ASH2L drives proliferation and sensitivity to bleomycin and other genotoxins in Hodgkin’s lymphoma and testicular cancer cells

**DOI:** 10.1101/2020.08.12.248559

**Authors:** Daniel Constantin, Christian Widmann

## Abstract

It is of clinical importance to identify biomarkers predicting the efficacy of DNA damaging drugs (genotoxins) so that non-responders are not unduly exposed to the deleterious effects of otherwise inefficient drugs. Using a whole genome CRISPR/Cas9 gene knockout approach we have identified that low levels of ASH2L cause resistance to genotoxins. ASH2L is a core component of the H3K4 methyl transferase complex. We show that ASH2L absence decreases cell proliferation and favors DNA repair upon genotoxin exposure. The cell models we have used are derived from cancers currently treated either partially (Hodgkin’s lymphoma), or entirely (testicular cancer) with genotoxins. For such cancers, ASH2L levels could be used as a biomarker to predict the response to genotoxins. Our data also indicate that patients with low ASH2L expressing tumors do not develop resistance to ATR inhibitors. In these patients, such inhibitors may represent an alternative treatment to DNA damaging drugs.

## Background

Cancer is the second leading cause of death worldwide, only surpassed by cardiovascular disease (1). In high income countries however, cancer is responsible for nearly twice as many deaths as cardiovascular disease, with a global trend towards cancer becoming the leading cause of death worldwide (2). One of the main anticancer therapeutic avenues are represented by DNA damaging agents (also called genotoxins) (3). The anti-cancer properties of these drugs were discovered in the 1940s. Genotoxins became very popular in the 1960s and 1970s (4). To this day, DNA damage causing agents are the largest class of anticancer drugs. They are used to treat lung, breast, bladder, stomach, ovarian, testicular, and other cancers (5). These genotoxic agents work by activating an intricate series of signaling pathways that, taken together, have been termed the DNA damage response (DDR). In eukaryotes, the DDR involves a tightly regulated series of transcriptional, post-transcriptional and post-translational events that have evolved to prevent transmission of damaged DNA to daughter cells. Initially the cell cycle is stopped and the DNA repair machinery is recruited to DNA lesions. If the extent of DNA damage surpasses the cell’s DNA repair abilities, additional pathways are activated that lead to cell death via mitotic catastrophe, apoptosis or necrosis, the ultimate outcome that is expected from genotoxins (6–8). Although these types of chemotherapy agents are used in many standard therapy regimens that represent the gold standard of care at the moment, they are also highly unspecific and almost always display systemic side effects that can be life threatening (9).

Here we have focused our initial investigation on a DNA damaging drug called bleomycin. This drug was discovered in the 1960s and FDA approved in 1973. It became very popular in oncology clinics due to its anti-tumor activity and low myelosuppressive properties (10, 11). Bleomycin is currently part of the gold standard therapy for Hodgkin’s lymphoma (HL), testicular cancer (TC), germ cell cancers and others but its mechanism of action is still not fully understood (12, 13). Akin to other unspecific chemotherapy drugs, bleomycin has severe side effects, mainly pulmonary toxicity that can be lethal on its own. Up to 46% of patients treated with bleomycin containing therapies develop pulmonary complications (14). These complications can lead to the patient’s death in 1-4% of cases (15–18). We focused on this drug because of a knowledge gap between the wide use of bleomycin in the clinic and the limited information about the genes implicated in the sensitivity or resistance to this compound. Here we show that HL and TC cancer cells depleted of ASH2L (Absent, Small, or Homeotic-Like 2) are more resistant to bleomycin and other DNA damaging agents. ASH2L is part of protein complexes responsible for the tri-methylation of histone 3 at lysine 4 (H3K4me3). These complexes are comprised of a catalytic subunit (either MLL1, MLL2, MLL3, MLL4, hSET1A or hSET1B) bound to the integral core members of the complex (ASH2L, RbBP5, and DPY30) (19). H3K4me3 is found mainly at promoters of transcriptionally active genes. Interestingly, bleomycin resistance in ASH2L knockdown cells does not seem to be mediated through changes in the gene expression, but rather through a global decrease in H3K4me3 levels that prime the chromatin for DNA repair. Within the context of personalized cancer therapy, decreased ASH2L levels could be used as a biomarker for predicting a poor response to DNA damaging agent-based therapy.

## Results

We conducted a whole genome CRISPR/Cas9 gene knockout screen (20, 21) in the presence or in the absence of bleomycin in L1236 Hodgkin’s lymphoma cells. These cells are derived from a type of cancer that is currently treated with bleomycin-containing chemotherapy. MaGECK algorithm (22) analysis of our CRISPR/Cas9 screen showed that ASH2L-depleted cells were enriched in the bleomycin-treated population (Figure 1A). All 6 sgRNAs targeting ASH2L present in our library were enriched in the population of cells treated for 10 days with bleomycin when compared to the untreated population cultured for the same period of time (Figure S1A). We used an independent genetic approach based on small hairpin RNAs (shRNAs) to investigate the effect of ASH2L depletion on genotoxin-resistance. The shRNAs-mediated decrease in ASH2L protein levels in L1236 cells was accompanied by H3K4 methylation reduction (Figure 1B). ASH2L knockdown in L1236 cells resulted in increased survival in response to bleomycin and to another genotoxin, etoposide, a topoisomerase II inhibitor (Figure 1C-E).

**Figure 1.**
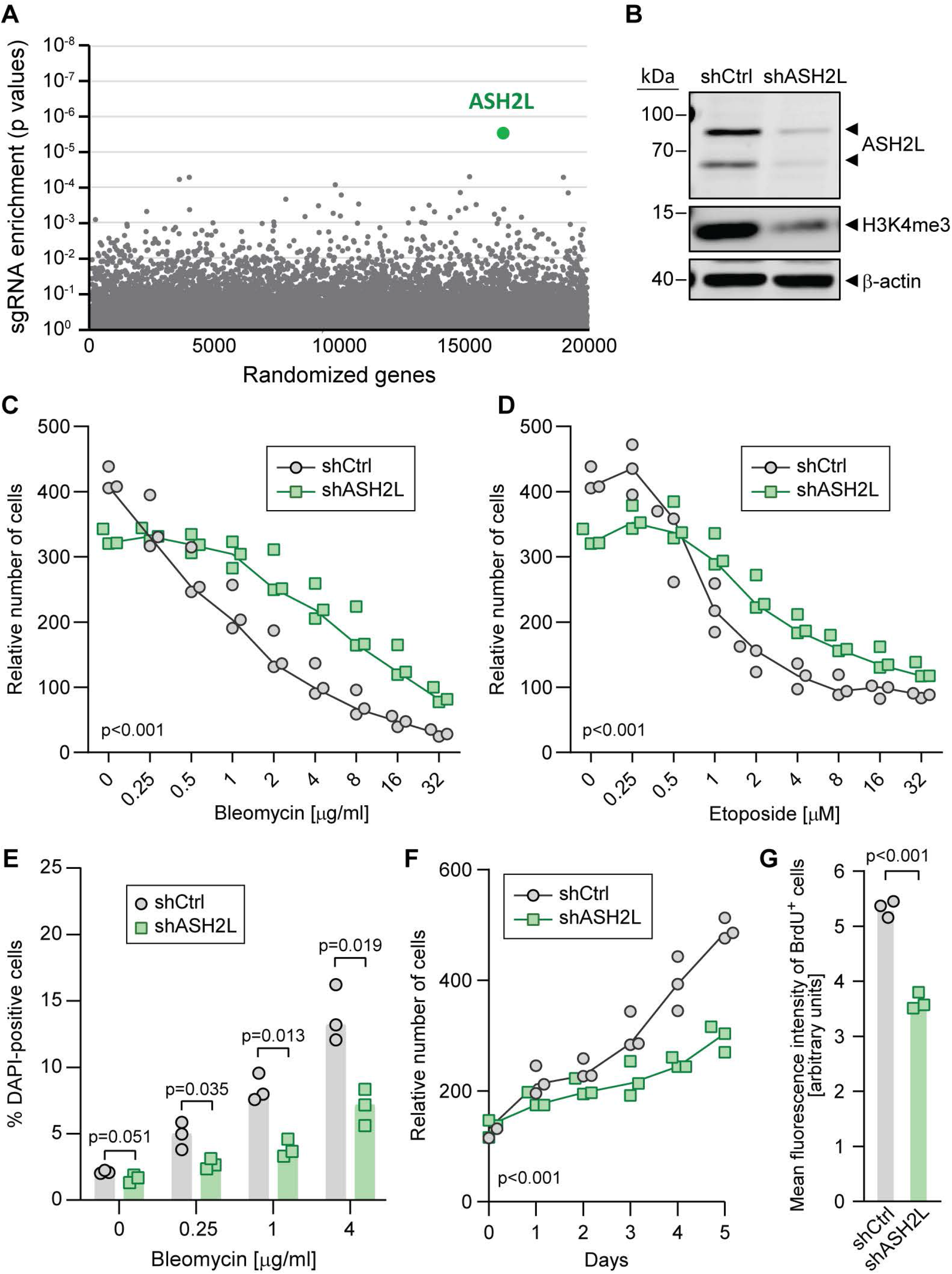
ASH2L depletion leads to resistance to genotoxins in Hodgkin’s lymphoma cells. **A.** MaGECK-based sgRNA enrichment analysis of the CRISPR/Cas9 whole genome knockout screen results. The graph depicts the p value associated with the observed difference in sgRNA abundance between untreated cells and cells treated with bleomycin (250 ng/ml) for 10 days. **B.** Western blot depicting shRNA-mediated knockdown of ASH2L and H3K4me3 levels. The arrowheads indicate the 2 ASH2L splice variants expressed in most tissues (75). **C-D**. Control (shCtrl) and ASH2L knockdown (shASH2L) L1236 cells were treated with increasing concentrations of bleomycin (panel C) or etoposide (panel D) for 3 days. The relative numbers of cells in the wells were estimated by Presto-Blue assays. **E.** ASH2L knockdown and control cells were treated with the indicated concentrations of bleomycin for 72 hours. The dead cells were stained with DAPI and then analyzed by flow cytometry. **F**. ASH2L knockdown and control cells were plated in 96 well plates and the relative number of cells was analyzed each day during 5 days. **G**. Flow cytometry analysis of the DNA synthesis rate, measured by BrdU pulse labelling, in control (shCtrl) and ASH2L knockdown (shASH2L) cells.

In the absence of bleomycin exposure, the mere culture for 10 days of the cell population expressing the sgRNA library led to a decrease in abundance of ASH2L-targeting sgRNAs (Figure S1B). ASH2L-targeting sgRNAs were among the top most depleted ones following a 10-day culture period (Figure S1C). This indicates that, in untreated cells, the absence of ASH2L decreases the ability of the cell population to expand, for example because of a reduced proliferation potential. A decrease in cell proliferation upon ASH2L depletion was indeed previously reported (23). We could reproduce the negative impact of ASH2L loss on population growth using shRNA-mediated ASH2L knock-down (Figure 1F). Cells lacking ASH2L had a decreased proliferation potential (Figure 1G), indicating that the reduced population growth upon ASH2L silencing is, at least partly, a consequence of decreased proliferation. Interestingly, none of the other genes that were detected as important for optimal population growth (e.g. PFN1, PPP4C, DUT or RPLP0) (Figure S1C) had the corresponding sgRNAs significantly enriched in the bleomycin-treated population (Table S1, first data sheet). This indicates that decreased proliferation is not the main mechanism driving the observed resistance to bleomycin upon ASH2L depletion.

One possibility that could explain the observed bleomycin-resistance phenotype is that a decrease in the H3K4me3 mark, which labels transcriptionally “open” chromatin, results in higher proportion of heterochromatin within the nucleus. This would make it more difficult for bleomycin to access DNA and cause double strand breaks. However, the levels of H3K9me3 and H3K27me3, the main heterochromatin markers (24), were not changed upon ASH2L knockdown in L1236 cells (Figure S2A and B).

We also checked if over-expression of ASH2L had an impact on L1236 cell survival upon genotoxin exposure or on H3K4me3 levels but found none (Figure S2D).

As stated above, H3K4me3 modification, which mainly occurs on promoters, is considered a mark of transcriptionally active open chromatin (24). At first sight, it was surprising that a protein required for this modification ended up to be the top hit in a screen designed to identify proteins modulating bleomycin sensitivity. While most publications are focusing on H3K4me3 in the context of transcription, there is evidence that this mark influences the DNA repair ability of cells. Firstly, upon DNA break induction, histone lysine demethylases are recruited to damaged chromatin to remove the H3K4me3 mark (25–27). Secondly, Bayo *et al.* have elegantly shown that augmenting H3K4me3 levels in cancer cells decreases their ability to repair DNA breaks efficiently, rendering cells more sensitive to DNA damaging agents (28). These studies prompted us to investigate the cells’ ability to repair their DNA upon ASH2L depletion. We used specifically engineered U2OS cells that carry reporters for the two main DNA repair pathways in mammalian cells: the DR-GFP reporter, developed by Maria Jasin (29), which measures repair through homologous recombination (HR), and the EJ5-GFP reporter, developed by Jeremy stark (30), which measures repair through non-homologous end joining (NHEJ). Figure 2A depicts how the reporters record these two types of DNA repair. ASH2L knockdown and reduction in H3K4me3 levels was achieved by transfecting the two reporter cell lines with a pool of 30 ASH2L-directed siRNAs (Figure 2B). Upon induction of DNA damage by I-SceI expression in the reporter cells lines, we observed a slight but significant increase in both homologous recombination and NHEJ repair efficiencies upon ASH2L silencing (Figure 2C). In accordance with the hypothesis that ASH2L is detrimental to DNA repair, endogenous ASH2L proteins were found to be excluded from areas of laser-damaged DNA (Figure 2D and Figure S2C).

**Figure 2.**
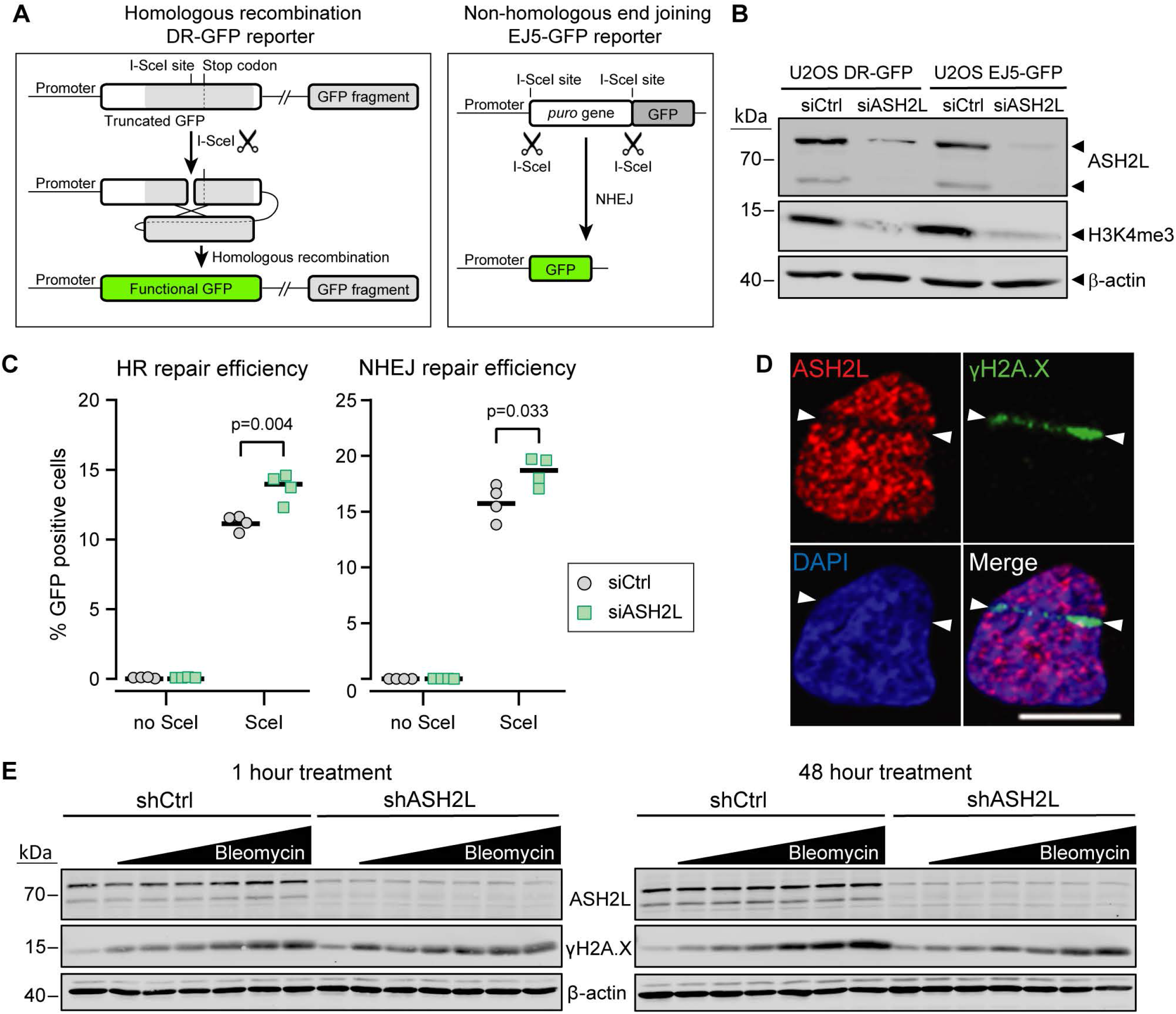
ASH2L depletion facilitates DNA repair. **A.** Schematic representation (not to scale) of the DR-GFP reporter (left) and EJ5GFP reporter (right). The DR-GFP construct bears a cDNA encoding the full-length GFP that contains an I-SceI restriction site disrupting the reading frame and leading to a premature stop codon. This cDNA is followed by a 5’- and 3’-truncated GFP fragment that does not contain the I-SceI restriction site. The sequences of this fragment and the complementary region in the full-length cDNA are indicated by different shades of grey. When cells expressing this reporter are transduced with a lentivirus encoding the I-SceI enzyme, the reporter is cleaved at the I-SceI site. The ensuing DNA repair by homologous recombination removes the termination codon in the full-length cDNA allowing for the production of a functional GFP (green). The EJ5GFP bears a puromycin resistance gene flanked by two I-SceI restriction sites. This gene is followed by a GFP-encoding cDNA. In cells expressing this reporter, removal of the puromycin resistance gene is achieved by transduction with an I-SceI encoding lentivirus. The ensuing NHEJ-mediated repair brings the promoter and the GFP cDNA close together, allowing for GFP production. **B.** Western blot depicting siRNA-mediated ASH2L knockdown efficiency in the U2OS DR-GFP and U2OS EJ5GFP reporter cell lines and the resulting decrease in H3K4me3. **C.** U2OS DR-GFP and U2OS EJ5GFP cells were transfected with control siRNAs or ASH2L-specific siRNAs for 48 hours and then infected with an I-Sce-I endonucleasecarrying lentivirus. Forty-eight hours post infection, the percentage of GFP positive cells was quantitated by flow cytometry (4 independent experiments; the horizontal bards correspond to the medians). **D.** Live U2OS wild-type cells were micro-irradiated using a confocal microscope. The cells were then fixed and stained with the indicated antibodies. Nuclei were labelled with DAPI. White arrowheads mark the boundaries of the irradiation track. Scale bar: 10 μm. **E.** L1236 cells were treated with 0, 0.06, 0.12, 0.25, 0.5, 1 or 2 μg/ml of bleomycin. Samples were harvested 1 hour (left) and 48 hours (right) post-treatment. The cells were lysed and 7.5 μg of protein were analyzed by Western blotting using the indicated antibodies.

We then tested if increased DNA repair activity is also detected in L1236 cells subjected to the selection conditions used in the CRISPR/Cas9 screen. Control and ASH2L-knockdown cells were therefore incubated with increasing bleomycin concentrations for one hour or for 48 hours and the extent of DNA damage evaluated by measuring phosphorylated H2A.X (γH2A.X) levels (31). After one hour of bleomycin treatment, γH2A.X levels were similar between control and ASH2L knockdown cells, indicating similar DNA damage induction in the two cell types. However, after 48 hours, these levels increased more in control cells compared to ASH2L knockdown cells (Figure 2E), supporting the notion that, like in U2OS cells, depleting ASH2L in L1236 cells leads to increased DNA repair capability.

As stated above, ASH2L is part of a protein complex that is responsible for the methylation of histone 3 at lysine 4. This chromatin modification is generally present at the promoters of active genes (32). We investigated the possibility that knocking down ASH2L leads to a gene expression signature responsible for the promotion of DNA repair and resistance to genotoxic stress. There is some evidence that points in this direction as ASH2L interacts with p53 at the promoters of pro-apoptotic genes (33). Note however that the Hodgkin’s lymphoma cell line in which we performed our screen is TP53 null (34). We performed three independent transcriptomic analyses of control (shCtrl) and ASH2L knockdown (shASH2L) in L1236 cells. Upon setting a threshold of fold discovery rate (FDR) of 0.05 and fold change ≥2, we found 93 up-regulated and 228 down-regulated genes in cells in which ASH2L was silenced compared to the control cells. The differential expression analysis for 14’000 genes is provided in Table S2. The majority of the up-regulated genes were low expressed genes (Figure 3A). The top 40 up-regulated genes are shown in Figure 3B and the top 40 down-regulated genes are shown in Figure 3C. We performed a gene ontology (GO) analysis on the genes that were up- and down-regulated (FDR < 0.05 and fold change ≥ 2). The main enriched GO term when analyzing the genes down-regulated in the ASH2L knockdown cells was cell proliferation regulation (Figure 3D). This is in agreement with the observation that these cells grow slower (Figure 1F). No enriched GO terms were found when analyzing the genes that were up-regulated upon ASH2L knockdown. These data indicate that resistance to bleomycin observed upon ASH2L depletion is not a consequence of increased expression of genes involved in DNA repair.

**Figure 3.**
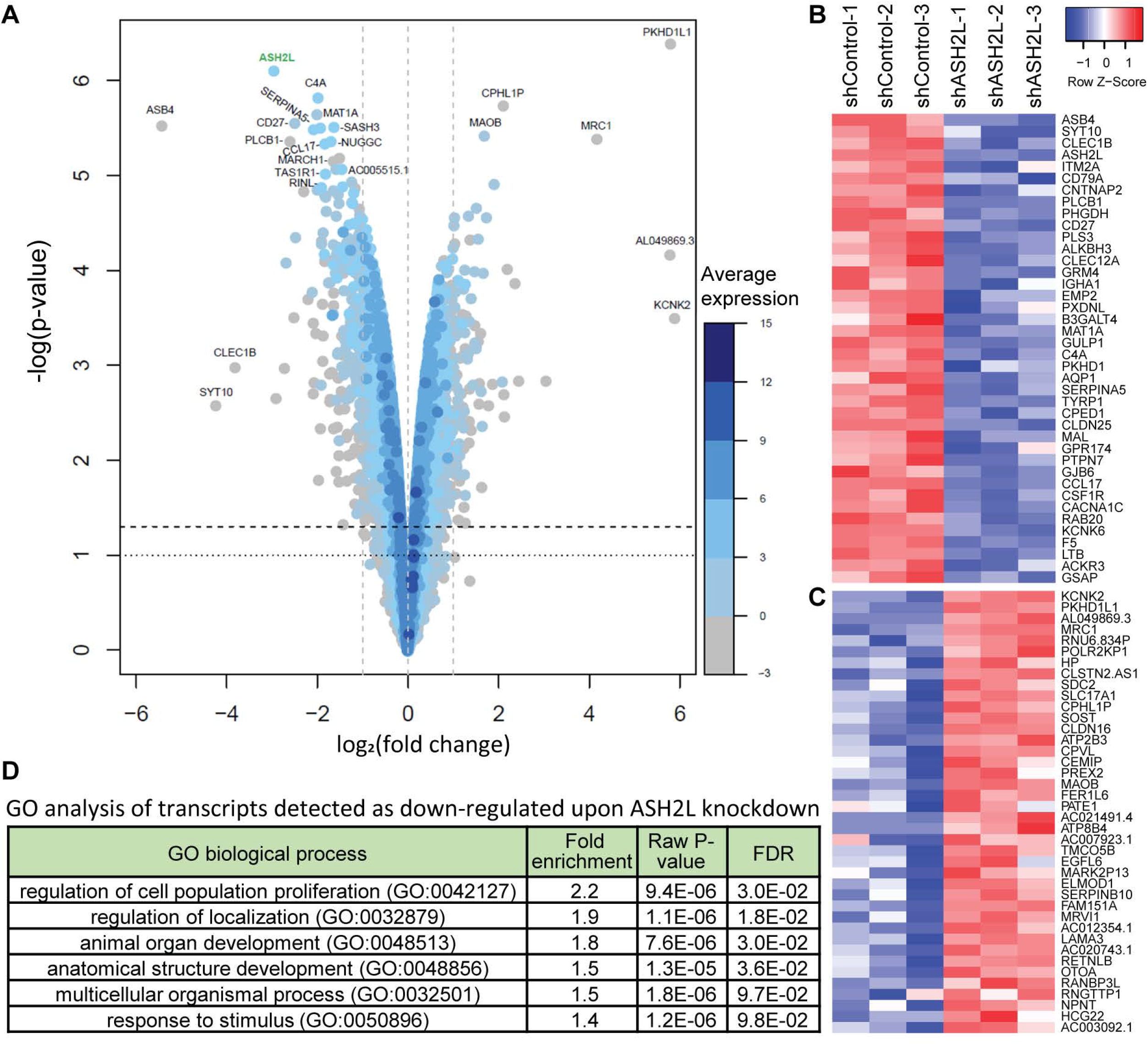
Transcriptome changes upon ASH2L depletion in L1236 cells. **A.** Volcano plot showing on the x-axis changes in gene expression levels and on the y-axis the p-values associated with these changes. The average expression of the genes is indicated by a color code (see scale bar on the right of the volcano plot). **B-C**. Heatmaps depicting, according to fold changes, the top 40 down-regulated (panel B) and up-regulated genes (panel C) with FDRs <0.05. Each column represents one independent experiment. **D**. Gene ontology (GO) analysis of genes down-regulated upon ASH2L silencing (FDR < 0.05 and fold change ≥ 2).

Since the resistance to genotoxins observed after ASH2L down-regulation does not seem to be regulated through changes in the transcriptome, we sought other possible explanations.

One possibility is that, upon DNA breaks being sensed, H3K4 demethylases are recruited to the lesion and remove this mark. Only then is the repair machinery recruited and the break repaired through HR or NHEJ. In recent years, a few studies have shown that hindering the cells’ ability to remove this mark at areas of DNA damage results in a decreased ability of cells to repair their DNA leading to subsequent sensitivity to genotoxins (25, 26, 28). Our results are in agreement with these observations. We extend these findings by showing, for the first time, increased DNA repair capabilities of cells with diminished H3K4me3 and/or ASH2L levels. One way to probe the hypothesis that low levels of H3K4me3 promote DNA repair is through overexpressing a histone lysine demethylase, such as KDM5A (35). Unfortunately, we were unable to obtain stable expression of KDM5A in cells using either retroviruses, inducible lentiviruses, or generation of stable cell lines through plasmid integration. We could obtain KDM5A expression by transfecting cells with KDM5A-expressing plasmids but for this we had to use five times the usual amounts of DNA during the transfection procedure (this “over” transfection was obtained using 5 μg of DNA per well of 12-well plates containing 10^5^ cells). This expression was functional, as evidenced by decreased levels of H3K4me3, but was very transient (Figure S3).

Our data indicate that cells with reduced H3K4me3 levels have a selective advantage compared to normal cells in repairing their DNA in response to genotoxin-induced DNA breaks. This is in line with earlier studies showing that a reduced ability to remove the H3K4me3 mark at areas of DNA damage hampers the recruitment of repair proteins, leading to subsequent sensitivity to genotoxins (25, 26, 28). We can therefore predict that H3K4me3 levels no longer influence cell survival when cells are prevented from turning on the signals that bring DNA repair proteins to sites of damaged DNA. This can be achieved through inhibition of ATM or ATR, the two main DNA lesion signaling kinases (36). In the absence of ATM or ATR, cells are blind to DNA damage and die from unrepaired DNA breaks, through mitotic catastrophe for example (37, 38). Figure 4 shows indeed that ASH2L-depleted cells have no survival benefit when incubated with ATM or ATR inhibitors (i.e. their IC_50_ values are similar to those of the control cells).

**Figure 4.**
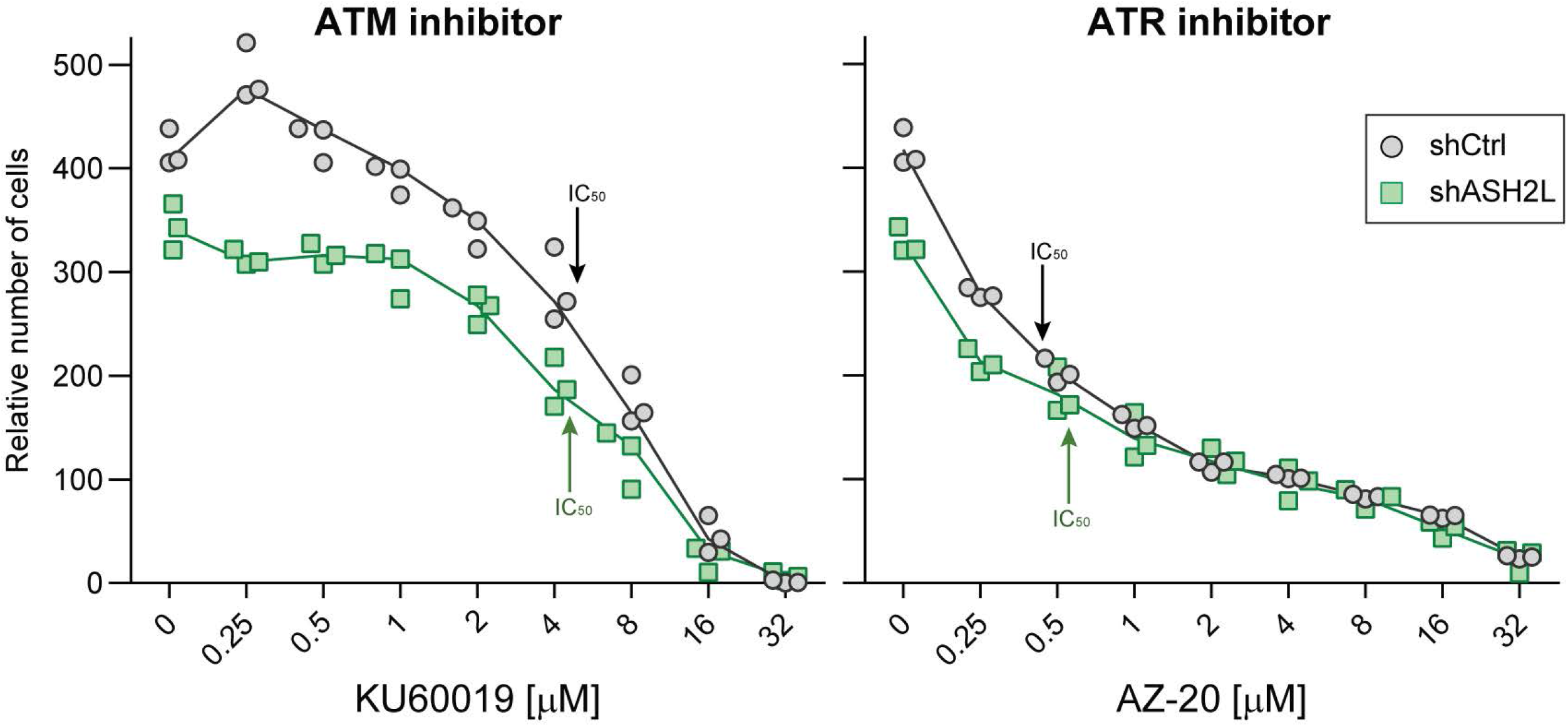
Effect of ASH2L depletion on ATM or ATR inhibition-induced death. Control (shCtrl) and ASH2L-silenced (shASH2L) L1236 cells were incubated with increasing concentrations of the KU60019 ATM inhibitor or the AZ-20 ATR inhibitor for 72 hours. The numbers of cells in the wells were quantitated using the Presto-Blue assay. Three independent experiments were performed. The curves go through the median values. The functionality of KU60019 is shown in Figure S4. IC_50_ values correspond to the drug concentrations that intersect the curves halfway between the highest and lowest y axis values.

We next focused on testicular cancer (TC) for which bleomycin is part of the standard therapy (bleomycin, etoposide, and cisplatin). TC represents around 2% of the total number of cancers in males and the most common malignancy in males aged 15 to 34 years. The incidence of testicular cancer has doubled in the past 40 years and continues to increase (39, 40). Due to the external location of the affected organs and the generally positive response to chemotherapy, TC has one of the highest cancer cure rates (approximately 95%) (41). Despite this high survival rate, there is a subset of patients with poor or even absence of response to standard chemotherapy (42). There are currently no consensus biomarkers for predicting the effectiveness of the standard chemotherapy used to treat TC (43). We transduced testicular cancer NT2D1 cells with either control shRNA or an shRNA targeting ASH2L and tested the effectiveness of the knockdown and the subsequent H3K4me3 decrease in these cells (Figure 5A). Strikingly, ASH2L knockdown cells were resistant to all 3 drugs used to treat this disease (Figure 5B). The observed resistance correlated with decreased DNA damage after 48 hours of drug exposure, indicating more efficient repair of damaged DNA in ASH2L-depleted cells (Figure 5C). Similarly to L1236 cells, overexpression of ASH2L in NT2D1 cells did not change their sensitivity to bleomycin (Figure S2E). Due to the observed genotoxin resistance in ASH2L knockdown cells, we checked if the available clinical data support the idea that ASH2L levels represent a prognostic marker in TC patients. Using the Tumor Cancer Genome Atlas (TCGA), available through the cBioportal website (44), we found no impact of ASH2L alterations on overall cancer survival. However, testicular cancer patients carrying alterations in the ASH2L gene were significantly more likely to relapse, indicating that the first line of treatment was not effective (Figure 5D). Lastly, we checked if there was any evidence of tumor samples that showed decreased levels of ASH2L gene expression. Interestingly, using TCGA data analyzed through the GEPIA software (45), we identified ASH2L as being overexpressed (at the mRNA level) in many TC samples compared to normal tissues. However, some TC tumor samples presented very low levels of ASH2L mRNA, even below the levels detected in normal tissues (Figure 5E). According to our hypothesis, the minority of patients with low ASH2L and/or H3K4me3 levels should be treated with non-DNA damage-based therapy in order for them not to suffer from the toxic side effects of non-effective genotoxic drugs.

**Figure 5.**
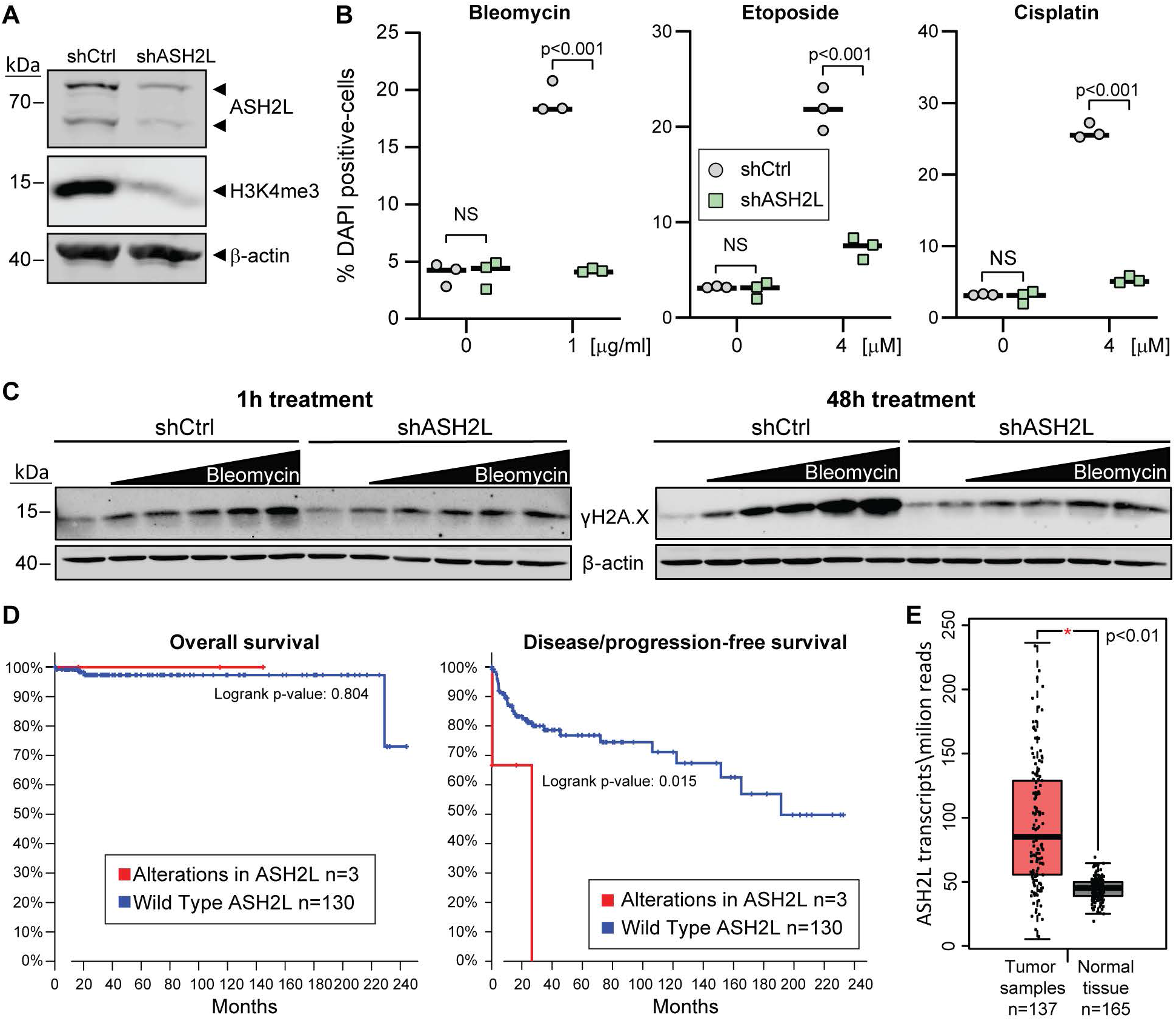
ASH2L depletion in the NT2D1 testicular cancer cell line leads to resistance to standard therapy. **A**. Western blot depicting shRNA-mediated ASH2L knockdown efficiency in NT2D1 TC cells. **B.** Control and ASH2L-silenced NT2D1 cells were treated with the indicated drugs for 72 hours. The percentage of dead cells of three independent experiments was quantitated by flow cytometry following DAPI staining. The horizontal bards correspond to the medians. C. Control and ASH2L-silenced NT2D1 cells were treated with 0, 0.12, 0.25, 0.5, 1 or 2 μg/ml of bleomycin. One set of cells was harvested at 1 hour and the second set at 48 hour post treatment. The cells were lysed and protein lysates analyzed by western blotting using the indicated antibodies. D. Testicular cancer patient survival (left) and relapse (right) graphs from the Tumor Cancer Genome Atlas (TCGA) database for patients with (red line) or without (blue line) alterations in the ASH2L gene. Data obtained through the cBioportal website. Cohort name: Testicular Germ Cell Tumors (TCGA, PanCancer Atlas). E. Graph comparing ASH2L transcript levels between testicular cancer samples and normal testicular tissue. The graph was generated through the GEPIA software. This tool requires a minimum fold change to be added to the analysis, we entered 1.74 (or log_2_FC=0.8), indicating that the ASH2L expression is significantly upregulated in cancer vs tumor samples and the fold change is at least 1.74. Dataset name: Testicular Germ Cell Tumors (TGCT).

## Discussion

The idea that cancer genome data can be used in order to predict the response to therapy of an individual, and offer potential guidelines to an optimal type of treatment, has been postulated since the first sequencing of the human genome (46). This concept is supported by recent meta-analyses of biomarker-based approaches in clinical oncology that have shown significant improvement in efficacy and overall death reduction caused by chemotherapy toxicity when a personalized approach is used compared to a non-personalized one (47, 48). Clinical oncologists generally agree that there is an urgent need for new biomarkers to predict the sensitivity or resistance to a particular treatment as this is a major limiting factor for contemporary precision medicine (49). The results presented here indicate that patients with low levels of ASH2L or H3K4me3 are more likely to relapse when treated with DNA damaging chemotherapy.

Although the overwhelming majority of studies investigating H3K4me3 modification do so from the perspective of transcription (19, 24, 50), some recent studies have focused on this modification in the context of DNA damage. Enzymes responsible for removing this mark, mainly members of the KDM5 family of proteins, are recruited to damaged DNA. Knockdown of these proteins decreases the ability of cells to repair their damaged DNA (25–27). Similarly, small molecule inhibitors of KDM5 enzymes increase the levels of H3K4me3 chromatin modification which in turn leads to impaired DNA repair through both homologous recombination and non-homologous end joining (28, 51). Our data are in agreement with these observations. We show that reducing the levels of H3K4me3 leads to increase in both of these repair pathways (Figure S5). In accordance with this, we show that ASH2L, a core component of the complex responsible for adding this chromatin mark, is excluded from areas of damaged DNA. Although decreased levels of H3K4me3 have previously been observed in areas of laser damaged DNA (25), it was not clear if this is only due to the recruitment of KDM5 enzymes or exclusion of the methylation complexes. Our data indicates that both scenarios occur simultaneously. In accordance with this model, Liu *et al.* have used colorectal cancer tissue samples to show a correlation between WDR82 protein levels, H3K4me3, and clinical outcome (52). WDR82 is a non-core member of the ASH2L-containing H3K4 trimethylation complex. Interestingly, patients with low H3K4me3 levels showed less aggressive tumors indicating low proliferation, but had a decreased overall survival compared with the high-expression group. One of the main avenues of treatment for colorectal cancer is radiation therapy, which induces double-stranded DNA damage. These observations are in line with our data obtained in Hodgkin’s lymphoma and testicular cancer cells.

There are studies, however, that indicate the exact opposite role of H3K4me3 in cancer progression. Decreased levels of H3K4 trimethylation levels have been suggested to be beneficial in terms of therapeutic outcome in pediatric ependymomas (53) and hepatocellular carcinoma (54). Further investigations will elucidate whether these discrepancies are due to differences between cancers derived from different tissues, the type of treatment used for these malignancies, or other factors.

Initially, it was surprising to us that no other core members of the complex responsible for H3K4 trimethylation were detected alongside ASH2L in our screen. However, both WDR5 and DPY30 depletion display an opposite phenotype as the one observed upon ASH2L knockdown. Down-regulation of WDR5 in colorectal cancer induces DNA damage and chemosensitivity (55), and knockdown of DPY30 increases the levels of endogenous DNA damage and decrease the DNA repair ability of hematopoietic stem cells (56). What seems to stand out about ASH2L is that it is the only core member of the methyl-transferase complexes that is required for the addition of the third methyl group. Unlike other core members of these complexes, knockdown of ASH2L does not affect the levels of H3K4me1 or H3K4me2 (57, 58). We currently lack the tools to fully distinguish the roles of each individual methyl group on histone 3 lysine 4. Moreover, Howe *et. al* argue that H3K4me3 is not required for, but more a consequence of efficient transcription, and that its actual function remains enigmatic (57). Although we could not fully disentangle the relationship between ASH2L, H3K4me3, and resistance to DNA damaging agents, the results presented here clearly indicate that low levels of ASH2L in cancer cells could be used to predict genotoxin resistance but not resistance to ATR inhibitors, which are in clinical trials at the moment (59).

## Conclusion

ASH2L was initially regarded as an oncogene (23). Our study indicates that it can also act as a tumor suppressor as its knockdown increases the resistance of tumor cell lines treated with various genotoxins (bleomycin, etoposide, cisplatin). However, ASH2L knock-down does not confer resistance to ATR or ATM inhibitors. Patients with low levels of ASH2L or H3K4me3 may more likely relapse when treated with DNA damaging chemotherapy but they should remain responsive to other anti-cancer drugs such as ATM or ATR inhibitors. ASH2L or H3K4me3 levels could prove useful as biomarkers in the clinic.

## Abbreviations

ASH2L: Absent, Small, or Homeotic-Like;
ATM: ataxia-telangiectasia mutated serine/threonine kinase;
ATR: ataxia telangiectasia- and Rad3-related serine/threonine kinase;
CRISPR: clustered regularly interspaced short palindromic repeats;
Cas9: CRISPR associated protein 9;
DUT: deoxyuridine 5’-triphosphate nucleotidohydrolase;
FDR: false discovery rate;
H2A.X: H2A histone family member X;
H3K4: histone 3, lysine 4;
H3K9: histone 3, lysine 9;
H3K27: histone 3, lysine 27;
me3: trimethylation;
MLL: mixed lineage leukemia protein;
hSET1: human SET domain-containing protein 1;
KDM5A: lysine-specific demethylase 5A;
PFN1: profilin 1;
PPP4C: protein phosphatase 4 catalytic subunit;
RBBP5: Retinoblastoma-binding protein 5;
RPLP0: ribosomal protein lateral stalk subunit P0;
SET: Su(var)3-9, Enhancer-of-zeste and Trithorax;
sgRNA: single guide RNA;
shRNA: short hairpin RNA;
siRNA: small interfering RNA.

## Declarations

### Ethics approval and consent to participate

Not applicable.

### Consent for publication

Not applicable.

### Availability of data and materials

Raw data concerning the CRISPR/cas9 screen and the transcriptomic analyses are available in the excel spreadsheet provided in the supplemental information. Additional raw data can be obtained from the corresponding author upon request.

### Authors’ contribution

Conception and design of study: DC and CW

Acquisition of data: DC

Analysis and/or interpretation of data: DC and CW

Drafting the manuscript: DC and CW

Revising the manuscript and approval of the submitted version: DC and CW

## Acknowledgements

We thank Dr. Nadja Chevalier and Gilles Dubuis for the amplification of the GECKO V2 library and producing the lentivirus stock used in this work.

## Methods

### Cell lines

L1236 Hodgkin’s lymphoma cells were purchased from DSMZ (ref: ACC 530) and grown in RPMI media. NT2D1 cells were purchased from ATCC (ref: CRL-1973) and grown in DMEM media. U2OS cells were purchased from ATCC (ref: HTB-96). U2OS cells with stably integrated homologous recombination (DR-GFP) and non-homologous end joining (EJ5GFP) reporters were a kind gift from the laboratory of Jeremy Stark (City of Hope Comprehensive Cancer Center, CA 91010). U2OS cells were grown in DMEM media. All cell culture media was supplemented with 10% fetal bovine serum and the cell lines were grown in 5% CO_2_ and a humidified atmosphere.

### CRISPR/Cas9 whole genome unbiased gene knockout screen

L1236 cells stably expressing the Cas9 protein were generated by transducing L1236 cells with viral particles created with LentiCas9-Blast (see details below). The cells were selected with 5 μg/ml blasticidin for 7 days. The GeCKO v2 library was purchased from Addgene (2-plasmid system catalog number 1000000049), and amplified in bacteria as previously described (60) and following the recommendations of the library creators (61). The library was packaged into viral particles (see lentivirus production section below). Then, we used 205·10^6^ L1236 cells stably expressing the Cas9 protein for transduction with the GeCKO viral library. Cells were plated at a density of 3·10^6^ cells per well of 12-well plates (in 3 ml media) and immediately infected with concentrated virus at a multiplicity of infection of 0.4 to insure that most cells only expressed one sgRNA. Sixteen hours later, cells were trypsinized, seeded in T150 flasks (200’000 cells/ml; 40 ml per flask) and cultured in the presence of 1 μg/ml puromycin to get rid of the non-infected cells. Seven days later, 7·10^7^ cells were harvested (“Time 0” sample), 7·10^7^ cells were left to grow untreated for 10 days (and passaged as necessary), and 7·10^7^ cells were grown for 10 days in the presence of 250 ng/ml bleomycin. The bleomycin-containing medium was changed every other day. The genomic DNA of the cells was extracted (see section below) and used undiluted for PCR amplification using primers binding 5’ or 3’ of the sgRNA sequence integrated in the cell’s genomes (62). In a 50 μl reaction, 7 μg DNA was used as template, 0.25 μM final primer concentration, and 25 μl of NEBNext High Fidelity PCR Master 2X Mix (New Englarnd Biolabs, M0541L). The primer sequences and their description, along with the PCR program, are available in Table S3. PCR amplicons were gel extracted (260 base pair band) and sent for next generation sequencing at the University of Lausanne, Genomic Technologies Facility (https://wp.unil.ch/gtf). Differential sgRNA distribution analysis between the control and ASH2L silenced cells was performed using the MAGeCK algorithm (22).

### Genomic DNA extraction

Cells were washed once in cold PBS and resuspended in lysis buffer (1% SDS, 50 mM EDTA, 50 mM Tris, pH 8). For 3.5·10^7^ cells, we used 6 ml lysis buffer. Next, we added 30 μl of 20 mg/ml proteinase K (Roche 03 115 828 001). The samples were incubated overnight at 55 °C. Afterwards, 15 μl of 20 mg/ml RNase A (Roche 10109169001) was added to the samples that were then quickly vortexed and incubated at 37°C for 30 minutes, followed by a 10 minute incubation on ice. At this point, 2 ml of cold 7.5M ammonium acetate was added to the samples that were then vortexed for 15 s and centrifuged at 4’000 x *g* for 10 minutes. The supernatant was transferred to a new tube on which 6 ml of isopropanol (100%) was added. The tubes containing the samples were inverted 15-25 times and then centrifuged at 4000 x *g* for 10 minutes. The supernatant was gently discarded and the remaining pellet was washed once with 70% ethanol. The ethanol was removed after an additional 4000 x *g* centrifugation for 2 minutes and the pellet was air dried for 5 minutes. The DNA pellet was resuspended in 0.5 ml TE buffer (Sigma T9285) and incubated at 65°C for one hour. The tubes were left to cool at room temperature and then the DNA concentration was measured via Nano-Drop apparatus (Thermo Fisher).

### Western Blotting

Cells were harvested by trypsinization (attached cell lines) or centrifugation (L1236 cells) and washed once with cold PBS. The cell pellets were resuspended in cold RIPA buffer (20 mM Tris-Cl pH 8.0,1 mM EDTA, 1% Triton X-100, 1% sodium deoxycholate, 0.1% SDS, 150 mM NaCl) supplemented with cOmplete protease inhibitor (Roche 11836145001) and PhosStop (Roche 04906837001), for 30 min on ice with occasional mixing. The samples were sonicated for 5 seconds at 40% amplitude using a Vibra-Cell 75186 machine. The tubes were centrifuged at 14000g, 4°C for 10 minutes and the supernatant was used for protein quantification using the BCA method (Pierce 23225). Samples were mixed with 2X Laemmli buffer and incubated for 10 min at 95°C. Thirty μg of protein was loaded per lane on either 15%, 10% or 4-20% polyacrylamide gels. The nitrocellulose membranes were blocked for 30 min at RT with 5% bovine serum albumin (BSA) dissolved in TBST solution. Antibodies were diluted (1:2000) in TBST with 5% BSA, and incubated overnight at 4°C with gentle shaking. Membranes were washed 3 times with TBST and incubated with fluorochrome conjugated secondary antibodies (Invitrogen, A21109, A21057, or A32735), for one hour at room temperature. Membranes were then washed 3 times with TBST and scanned using a Lycor Odyssey system.

### Presto-Blue cell number assay

We used the Presto-Blue viability (Thermo A13262) reagent according to the manufacturer’s instructions. Briefly, L1236 cells were plated at 2·10^3^ cells per well of 96-well plates containing 100 μl of media per well. The cells were then treated with the indicated doses of drugs for 72 hours. Presto-Blue reagent was added to each well (10 μl/well) and the plates were incubated for 2 hours at 37°C and 5% CO_2_. The fluorescence was read at 560/590 nm (excitation/emission) using a Cytation-3 plate reader. The value of the blank samples (media with no cells but with Presto-Blue) was deducted from all wells and the remaining absorbance values were plotted in the graphs. For the proliferation assay shown in Figure 1F, control and ASH2L depleted L1236 cells were plated in 96-well plates at 1000 cells per well. Presto-Blue assay was performed each day, from day 0 to day 6.

### DAPI viability assay

Cells were plated in 12-well plates (4·10^4^ for NT2D1 cells, and 1·10^5^ for L1236 cells) and treated with genotoxins for the indicated amounts of time. Floating and attached cells were harvested and incubated directly (without washing) with DAPI (Thermo 62248) at a final concentration of 1μg/ml in the media they were grown in. After 10 minutes the cells were analysed on a CytoFlexS flow cytometer, on the KO525 channel. A gate was placed on the single cells, and subsequently on the DAPI-positive cells.

### 5-bromo-2’-deoxyuridine (BrdU) incorporation assay

L1236 cells were infected with control and ASH2L-specific shRNA-encoding viruses and grown in the presence of doxycycline for 4 days in order for the knockdown to occur (see the “shRNA-mediated knockdown of ASH2L” section below). Thirty minutes before harvesting, BrdU (10 μM final concentration) was added to the cell cultures. Unless otherwise stated, the next steps were performed at 4°C. Two million cells were harvested and washed with 1.5 ml PBS, and then resuspended in 300 μl PBS. Next, 700 μl 100% ethanol was added to the tubes under gentle vortexing to prevent clumping. The cells were then kept overnight at 4°C. The next day, the cells were washed once with 1.5 ml of PBS and then incubated in 500 μl 0.1 M HCl, 0.5% Triton X-100 in PBS for 10 min. The tubes were centrifuged at 170 x g for 5 minutes (we recommend a swing-out centrifuge for these samples, as the cells can stick to the sides of the tubes in regular angled centrifuges). The pellets were resuspended in 500 μl water, boiled for 10 minutes, and cooled down on ice for 10 minutes. Then, 1 ml of 0.5% Triton X-100 in PBS was added and the tubes were centrifuged as above. The pellets were resuspended in 50 μl PBS containing 1% BSA and 0.5% Tween and 1 μl of primary FITC-labeled anti-BrdU antibody added to each sample. The tubes were placed in the dark for one hour. Then, 1.45 ml of PBS was added and the cells were centrifuged (170 x g for 5 minutes), the pellets were resuspended in 500 μl PBS and analyzed by flow cytometry using the FITC channel of a FC-500 flow cytometer (Beckman Coulter). All manipulations were performed in 1.7 ml Eppendorf tubes. The data were analyzed as follows: a gate was placed on single cells, and a second gate (dependent of the first one) was placed on FITC positive (BrdU positive) cells. The geometric mean of all events in the BrdU positive gate was used to generate the graph in Figure 1G.

### DNA repair reporter assays

The U2OS cells with stably integrated homologous recombination (DR-GFP) and non-homologous end joining (EJ5-GFP) reporters were a kind gift from Jeremy Stark (City of Hope, Beckman Research Institute of the City of Hope, CA, 91010, USA). These cells were plated in 6-well plates (8·10^4^ cells/well) and immediately transfected with a pool of control siRNAs or a pool of siRNAs targeting ASH2L (see siRNA transfection protocol below). The next day, siRNA transfection was repeated. On the following day, the cells were transduced with a virus (multiplicity of infection of = 1) carrying an I-SceI endonuclease gene. The next day, media was changed and the cells were left to grow for another 48 hour period. The cells were then trypsinized and washed once with PBS. One quarter of the cells was used for western blot analysis to check knockdown efficiency. The rest of the cells was analyzed on a CytoFlexS flow cytometer, gaiting on GFP-positive single cells.

### RNA isolation and sequencing and analysis

L1236 harboring inducible shRNAs were plated at a 2·10^6^ cells per 10-cm plate density and stimulated 4 days with 1 μg/ml doxycycline. The cells were washed twice in cold PBS and total RNA was extracted using a High Pure RNA isolation kit (Roche 11828665001). Reverse transcribed RNA was used for sequencing by the University of Lausanne, Genomic Technologies Facility. Fastq files were demultiplexed and adapters were trimmed using Cutadapt (63). Fastq screen was used to remove ribosomal RNA sequences. STAR (64) was used to align the reads against the *Homo sapiens* GRCh38.92 genome. Differential gene expression was computed with limma (65) by fitting the 6 samples into a linear model and performing the comparison between shASH2L and shControl. As the experiment was performed in batches and samples are paired, pairing was corrected for in the linear model. Moderated t-test was used for the comparison between shCtrl and shASH2L samples. In order to correct for multiple testing, we used the Benjamini-Hochberg method (66) to calculate the false discovery rate (FDR, also known as adjusted p-value). Heatmaps were generated from log_2_ expression values, using the expression module of the heatmapper.ca online application. The Z score in the heat-maps is calculated by subtracting the mean of the row from each value and then dividing the resulting values by the SD (standard deviation) of the row. The gene ontology analysis was performed using the online Protein ANalysis THrough Evolutionary Relationships (PANTHER) classification system developed by Huaiyu *et al.* (67).

### Laser microirradiation

Wild-type U2OS cells were plated on glass bottom 35 mm culture dishes (MaTek P35G-1.5-14-C) at a density of 5·10^5^ cells per dish and incubated overnight. Ten minutes before transporting the cells to the confocal microscope, Live-Hoechst (Invitrogen H3570) was added to the cells at a final concentration of 1 μg/ml. This allows for the visualization of nuclei and also acts as a sensitizing agent. The live cells were microirradiated using the 405 nm X-Cite light source and the fluorescence recovery after photobleaching (FRAP) module of a Zeiss LSM 710 confocal microscope. Stripes within cells were irradiated using 100% of the laser power (5 iterations, pixel dwell time 50.1 μs). Fifteen minutes post irradiation, the cells were washed once with PBS and then incubated with CSK+R buffer (10 mM Pipes, pH 7.0, 100 mM NaCl, 300 mM sucrose, and 3 mM MgCl_2_, 0.5% Triton X-100, and 0.3 mg/ml RNAse A) two times for 3 minutes at room temperature to remove soluble proteins. The cells were then washed once with 2 ml PBS and fixed with 4% paraformaldehyde for 15 minutes. This was followed by two PBS washes and the addition of 2 ml of PBS containing 5% fetal bovine serum and 0.3% Triton X-100. The samples were co-incubated with anti-ASH2L and anti phospho-H2A.X antibodies diluted (according to their datasheet) in PBS containing 1% bovine serum albumin and 0.3% Triton X-100, overnight at 4°C. The next day, the plates were washed twice with 2 ml PBS and the secondary antibodies were added, diluted in the same buffer. One hour later, the plates were washed, and PBS containing 1 μg/ml Live-Hoechst was added for 10 minutes at room temperature. The cells were washed three times with PBS and glass coverslips were mounted using VectaShield (Vector laboratories H-1000-10), and sealed with nail polish. The next day, the dishes were imaged using the same confocal microscope that was used for the irradiation.

### Lentivirus production

Low passage HEK 293T cells were plated (1·10^6^ cells/plate; in 10 ml media) in 10 cm dishes, and left to adhere overnight. The cells were transfected using the calcium phosphate method (68), with 7.5 μg psPAX2, 2.5 μg pMD2.G, and 10 μg of specific cDNA or shRNA lentiviral-encoding plasmids. Briefly, chloroquine (Sigma C6628) was added to the media to a final concentration of 25 μM. In a sterile tube, the DNA and sterile water were mixed to a final volume of 450 μl. After the addition of 50 μl of 2.5 M CaCl_2_ solution, the samples were mixed and incubated 20 min at room temperature. Then, 500 μl of a 2X HEPES solution (NaCl 280 mM, KCl 10 mM, Na_2_HPO_4_ 1.5 mM, D-glucose 12 mM, HEPES 50 mM) were added and the tube contents were briefly mixed. One minute after the HEPES buffer was added, the contents of the tube were added dropwise to the cells. Eight hours later, the media was changed and the cells were grown for 48 more hours. Next, the media was collected and centrifuged at 2’500 rpm for 5 minutes to remove floating cells. The resulting supernatant containing the viral particles was aliquoted and stored at −80°C. Titration was performed using puromycin selection of infected cells. Unless specified otherwise, the minimal volume of viruses conferring 100% protection against puromycin was used in subsequent experiments.

### siRNA transfection

U2OS cells were plated in 1 ml of DMEM at 4·10^4^ per well in 6-well plates. The first round of siRNA transfection was performed right after the plating when the cells were still floating. The transfection mix was made as follows: 200 μl Opti-MEM (Thermo 11058021) were added to a sterile 1.7 ml Eppendorf tube, then 1.2 μl of siRNA (5 μM stock) was added, followed by 2 μl of Lipofectamine RNAi-MAX (Invitrogen 13778-075). The tube was gently mixed and incubated at room temperature for 15 minutes. This mixture was added dropwise in the well containing the cells. The next day the media was replaced with 1 ml of fresh media and the transfection was repeated one more time. The cells were analyzed 96 hours after the first transfection. The final siRNA concentration in the media was 5 nM. The pools of 30 siRNAs targeting ASH2L (ref# 9070), and the siRNA negative control (ref# si-C005-NEG001, a pool of 30 non-targeting siRNA sequences), were purchased from the siTOOLS Biotech company.

### shRNA-mediated knockdown of ASH2L

L1236 and NT2D1 cells were transduced with doxycycline-inducible shRNA carrying lentiviruses at an MOI of 0.5 (this was achieved with about 0.25 ml of non-concentrated viral supernatant added to 2 ml of media containing 10^6^ cells). We checked via western blotting that there was no ASH2L knockdown in the absence of doxycycline. For knockdown experiments, 2?10^6^ cells were plated in 10 cm^2^ plates, and cultured in the presence of 1 μg/ml of doxycycline for 4 days. Afterwards, the cells were re-counted and plated for experiments in 6, 12, or 96-well plates, depending on the type of experiment performed. The shRNA sequence targeting ASH2L was: 5’-TTTACCAAGAATACATCTC-3’ and the control shRNA sequence was: 5’-CACACAACATGTAAACCAGGGA-3’.

### cDNA transfection

NT2D1 cells were plated in 12-well plates, at 1 · 10^5^ cells per well, in 700 μl media, and left to adhere overnight. The next day the media was changed for 700 μl fresh media. In a sterile tube, 1 μg DNA was mixed with 100 μl Opti-MEM media, and 1 μl Lipofectamine 2000 (Invitrogen 11668-019) was added. The tube was gently mixed and incubated at room temperature for 30 minutes. The mixture was added dropwise in the wells. The media was changed the next day, and cells were analyzed 48 hours post-transfection.

### Antibodies

The antibody against ASH2L used for Western blots was from Bethyl (A300-489; lot 2) and the one used for immunocytochemistry was from Cell Signaling (CLS D93F6; lot 1). The antibodies against H3K4me3 (Ab8580; lot GR3264593-1), H3K9me3 (Ab176916; Lot GR2318257-2), H3K27me3 (Ab192185; lot GR3264827), and phospho-ATM at serine 1981 (Ab81292; lot GR217573-6) were purchased from Abcam. The antibody against phospho-H2A.X at serine 139 was acquired from Milipore (5636; lot 2554898), and the one against beta actin from Cell signaling (CLS4970; lot 14). Our anti FLAG antibody was purchased from Sigma (F1804-1MG; lot SLGB5673V). The FITC-labeled anti-BrdU antibody was purchased from Thermo Fisher Scientific (11-5071-42, lot 4315462)

### Chemicals

The following chemicals were purchased from Sigma-Aldrich: ATM inhibitor KU-60019 (SML1416-5MG), ATR inhibitor AZ-20 (SML1328-5MG), bleomycin (B8416), etoposide (E1383), doxycycline (D3072), cisplatin (P4394), 5-Bromo-2’-deoxyuridine (B5002. Puromycin was purchased from Thermo-Fisher (A1113802), blasticidin S was purchased from Carl Roth (CP14.2).

### Statistical analysis

For the dose response curves, the area under the curve (AUC) was calculated for each group (shCtrl and shASH2L) on a linear scale. Please note that the X axes in the figures are not displayed linearly in order to provide more clarity at low drug concentrations. The resulting AUC values and the standard errors were used to perform an unpaired t-test between the two groups. The “n” for the t-test was not 3. The “n” was equal to the degrees of freedom (df) of each curve. The df was calculated as follows: number of data points used for one curve (27) minus the number of concentrations (9) + 1 (because the t-test subtracts 1 automatically). All calculations were performed in GraphPad 8 software (La Jola, California, USA) according to AUC statistics module and validated by the University of Lausanne Biostatistics platform (https://wp.unil.ch/biostatistics/). The same analysis was done for the growth curve in Figure 1F. For Figures 1E and S2D, we performed unpaired t tests, and the p-values were corrected for multiple testing with the Benjamini and Hochberg method. For Figures 1G, 2C, 5B, and S2E, we performed two-tailed unpaired student’s t tests. At least 3 independent experiments were performed. Data are shown as median together with the individual measurements.

### Plasmids

LentiCas9-Blast (#849) is a third generation lentiviral vector expressing the human codon-optimized *S. pyogenes* Cas9 protein (tagged at the C-terminus with FLAG) and blasticidin (both genes driven by the EFS-NS promoter). This plasmid was a gift from Feng Zhang (Addgene plasmid # 52962; http://n2t.net/addgene:52962; RRID:Addgene_52962) (69).

The pcDNA3 plasmid (#1) is a mammalian, CMV-driven, expression vector.

The hASH2L.dn3 (#1055) pcDNA3-based plasmid encodes the human ASH2L 80 kDa isoform (NM_001105214.2), the most commonly expressed isoform in cells and tissues. This plasmid was a gift from Kai Ge (Addgene plasmid #15548; http://n2t.net/addgene: 15548; RRID:Addgene_15548) (70).

The pLentiPURO-tetO-V5-6xHis (#980) is a lentiviral backbone for mammalian expression of cDNAs. This plasmid was a gift from Ie-Ming Shih (Addgene plasmid # 39481; http://n2t.net/addgene:39481; RRID:Addgene_39481) (71).

The hASH2L.plp (#1052) is a hASH2L-encoding lentiviral vector. The plasmid was made by cutting out the hASH2L insert from hASH2L.dn3 (mentioned above, #1055) using BamHI and XhoI and ligating it into the receiving lentiviral vector (pLentiPuro) opened with the same enzymes.

The HA-FLAG-hKDM5A.dn3 (#1048) was a gift from William Kaelin (Addgene plasmid # 14799; http://n2t.net/addgene:14799; RRID:Addgene_14799) (72). It expresses N-terminally HA- and FLAG-tagged human lysine demethylase KDM5A in the pcDNA3 backbone.

HA-NLS-scSceI-IRES-BFP.pCVL (#1053) plasmid expresses the HA-tagged I-SceI-encoding sequence followed by an IRES and the coding sequence for the Blue Fluorescent Protein (BFP). There is a nuclear localization sequence (NLS) inserted between the HA tag and I-SceI. This construct was a gift from Andrew Scharenberg (Addgene plasmid #45574 http://n2t.net/addgene:45574; RRID:Addgene_45574) (73).

The shCtrl.lti (#1056) and shASH2L.lti (#1057) were purchased from Horizon Discovery (VSC11655 and V3SH11252-225078488, respectively). They code for shRNA sequences (described above under the “shRNA-mediated knockdown of ASH2L” section) and Turbo-RFP that are both under a doxycycline-inducible promoter.

The packaging vector psPAX2 (#842) and the envelope vector pMD2.G (#554) were gifts from Didier Trono (Addgene plasmids #12260 and #12259, respectively) (74).

## Legends to supplementary figures

**Figure S1.**
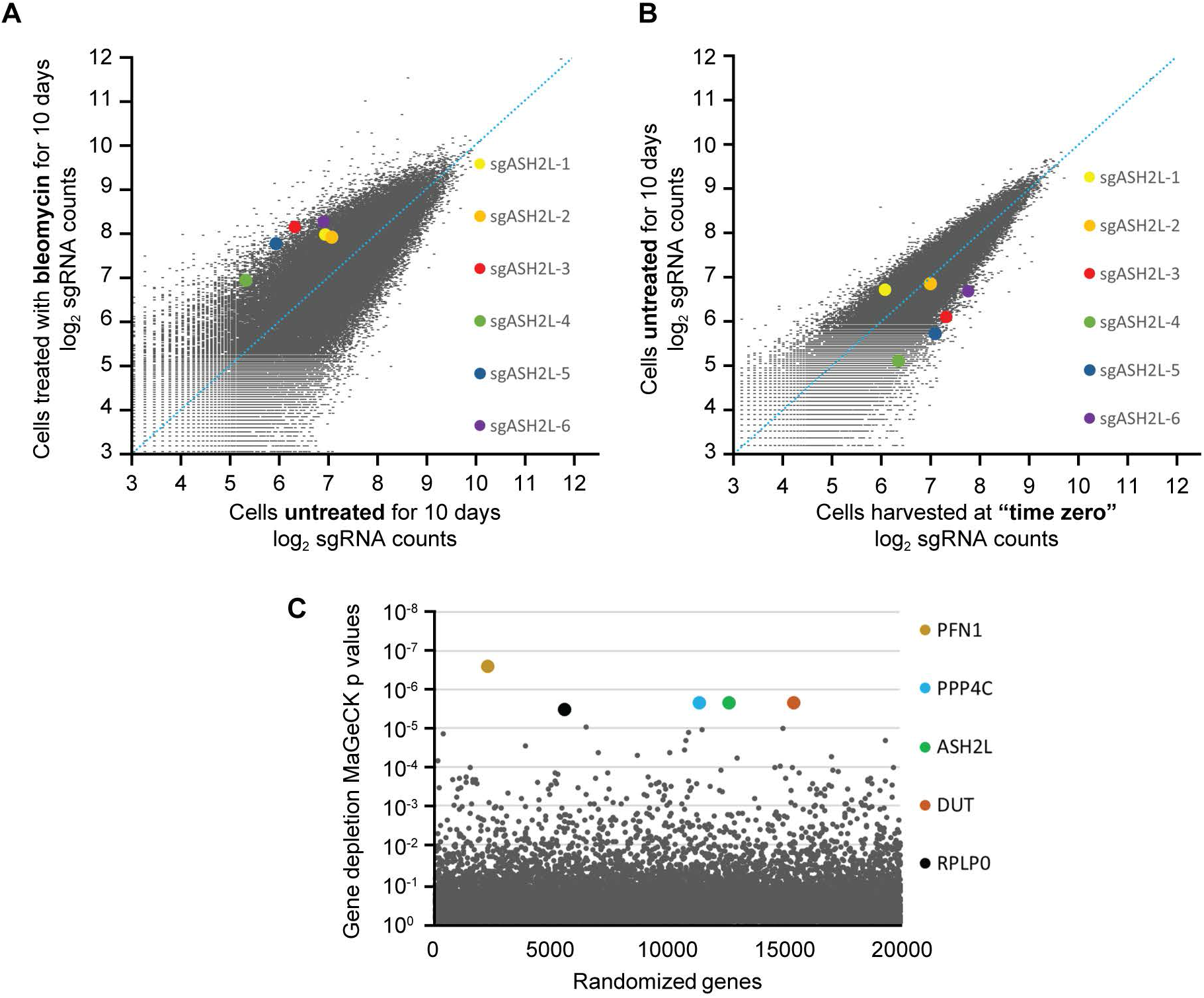
CRISPR/Cas9 whole genome knockout screen results in L1236 cells. **A.** Graph showing the prevalence of individual sgRNAs targeting ASH2L in DNA extracted from untreated (x axis) and bleomycin treated cells (y axis). **B.** Graph showing the prevalence of individual sgRNAs targeting ASH2L in DNA extracted from cells harvested at “time zero” (x axis) and cells left to grow untreated for 10 days (y axis). **C.** MaGeCK-based gene depletion analysis of the CRISPR/Cas9 screen results highlighting in color the top 5 depleted genes after 10 days of growth in normal medium (the analysis compared the sgRNA expression between cells harvested at “time zero” and cells left to grow untreated for 10 days).

**Figure S2.**
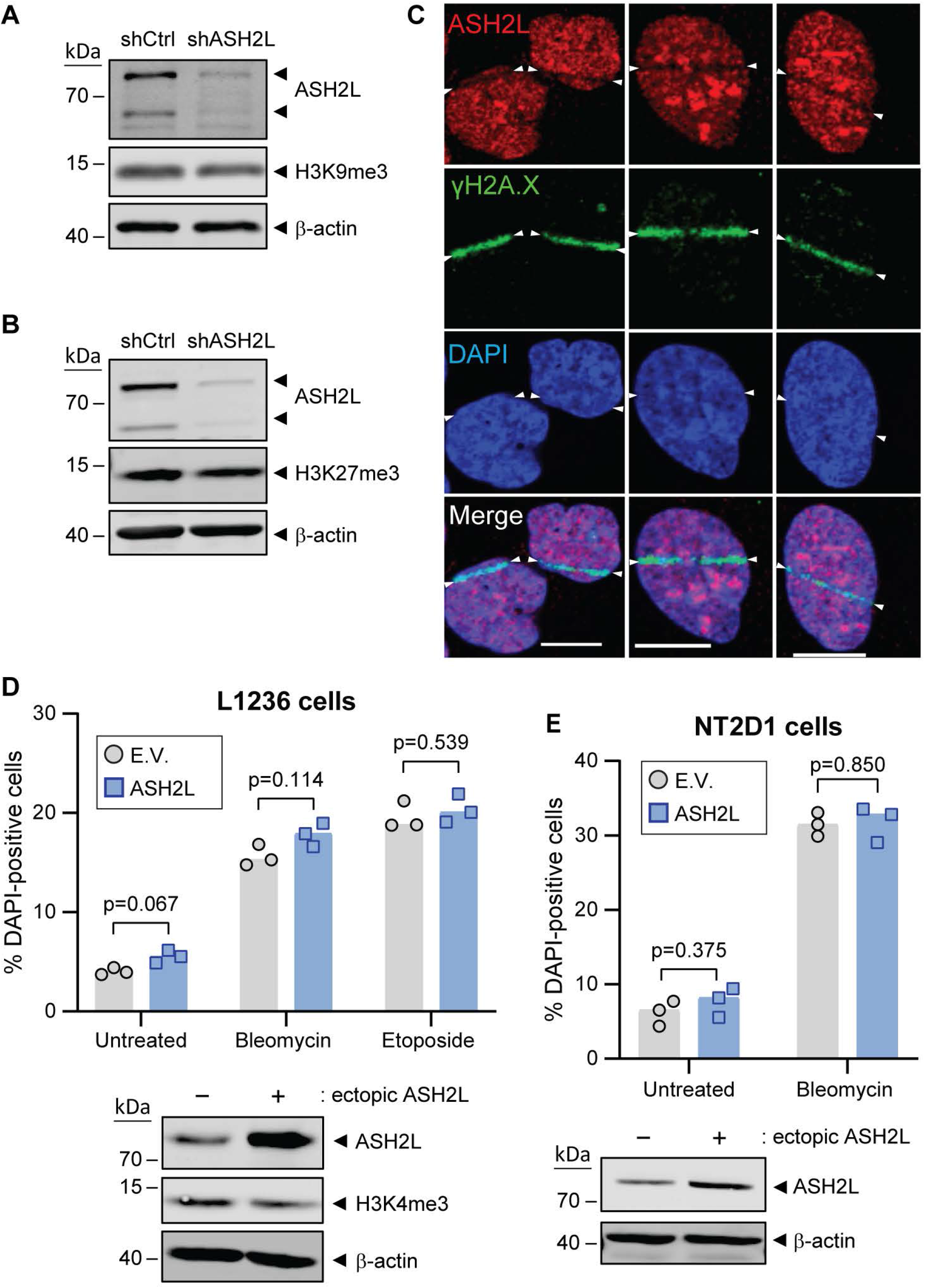
The impact of ASH2L knockdown on closed chromatin markers, localization of endogenous ASH2L at DSBs, and effect of ASH2L overexpression on genotoxic stress response. **A-B.** Western blot depicting shRNA-mediated knockdown of ASH2L and the corresponding levels of H3K9me3 (panel A) and H3K27me3 (panel B) in L1236 cells. **C.** Wild-type U2OS cells were laser-micro-irradiated and then immuno-stained with the indicated antibodies. DAPI was used to stain the nuclei. Three examples are shown. The white arrowheads mark the boundaries of the irradiated stripes in the nuclei. Scale bars: 10 μm. **D.** Top, the effects of lentiviral mediated ASH2L overexpression, compared to empty virus (E.V.) transduced L1236 cells, challenged for 72 hours with 4 μg/ml bleomycin, 4 μM etoposide, or left untreated. Bottom, western blot showing the levels of ASH2L overexpression and corresponding H3K4me3 levels. **E.** Top, NT2D1 cells transiently transfected with either pcDNA3 empty vector (E.V.) or hASH2L.dn3 (ASH2L), challenged for 72 hours with 4 μg/ml bleomycin, or left untreated. Bottom, western blots depicting the levels of ASH2L in these cells.

**Figure S3.**
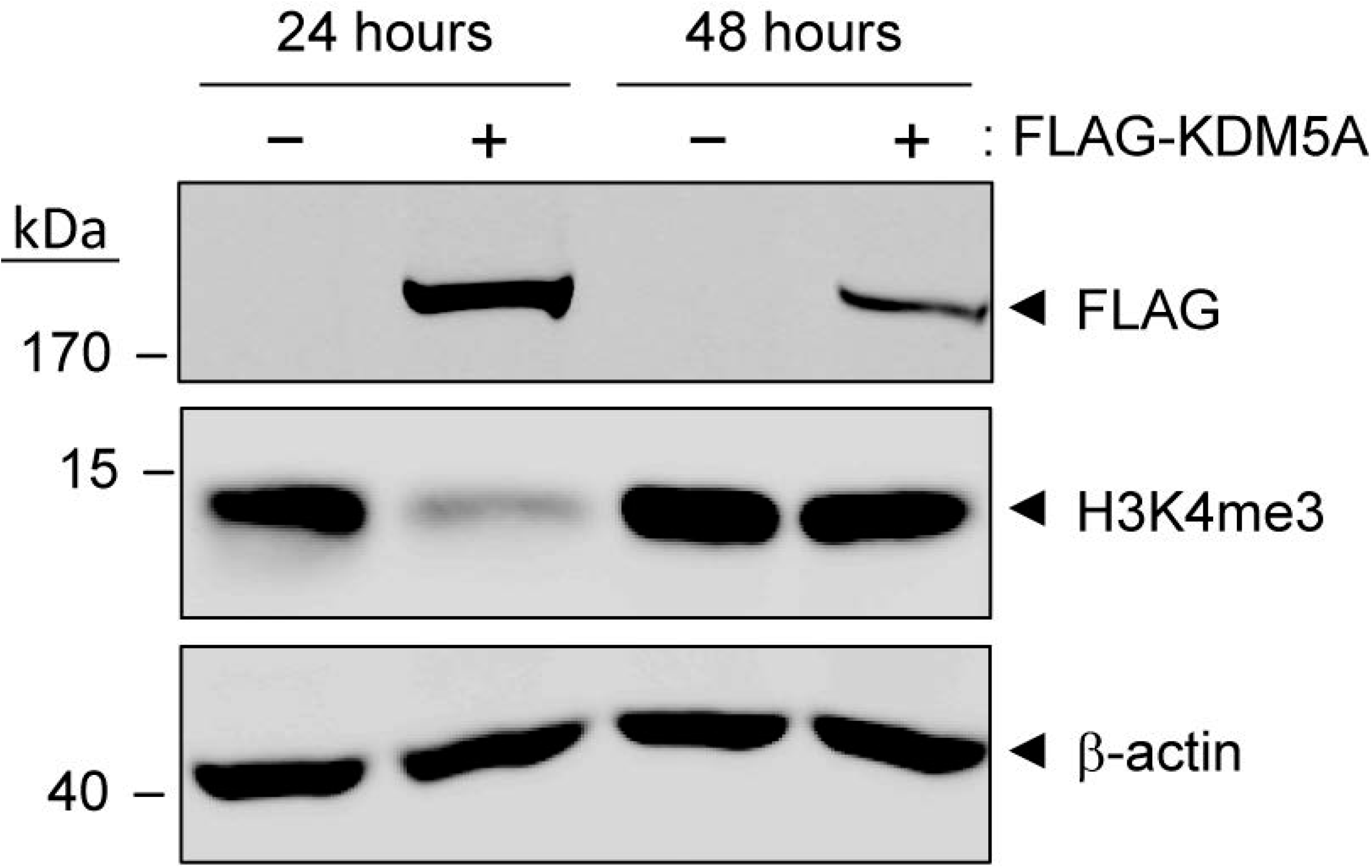
Transient nature of KDM5A ectopic expression. HEK293T cells were transfected with an empty pcDNA3 vector (-) or HA-FLAG-hKDM5A.dn3 (+). The cells were lysed at the indicated post-transfection times and the lysates analyzed by western blotting using the indicated antibodies.

**Figure S4.**
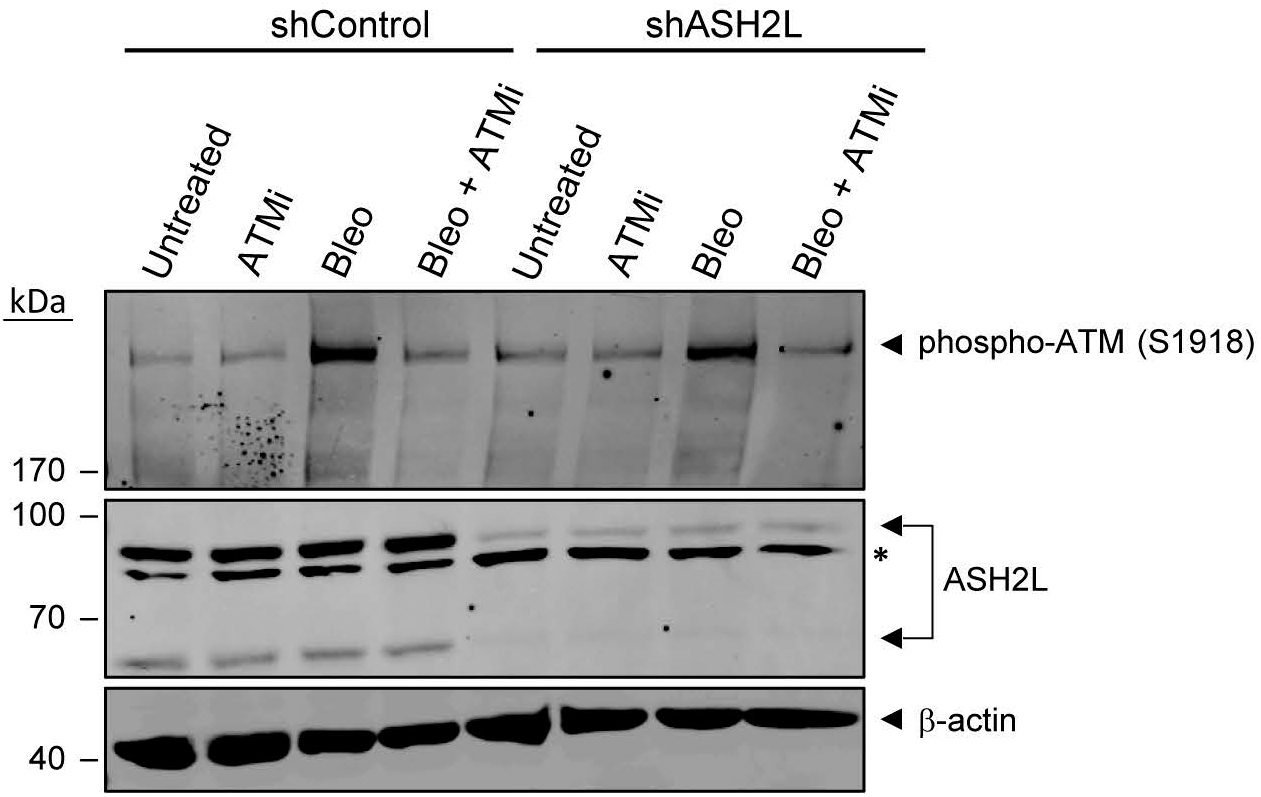
ATM inhibitor functionality test. L1236 control (shCtrl) and ASH2L-depleted (shASH2L) cells were pre-incubated for one hour with 2.5 μM of the KU-060019 ATM inhibitor (ATMi) before the addition, or not, of bleomycin (2 μg/ml; Bleo) for 2 more hours (still in the presence of KU-060019). The cells were then lysed and the lysates analyzed by western blotting using the indicated antibodies. Phosphorylation of ATM at position S1918 is indicative of an active kinase. The asterisk points to a non-specific band.

**Figure S5.**
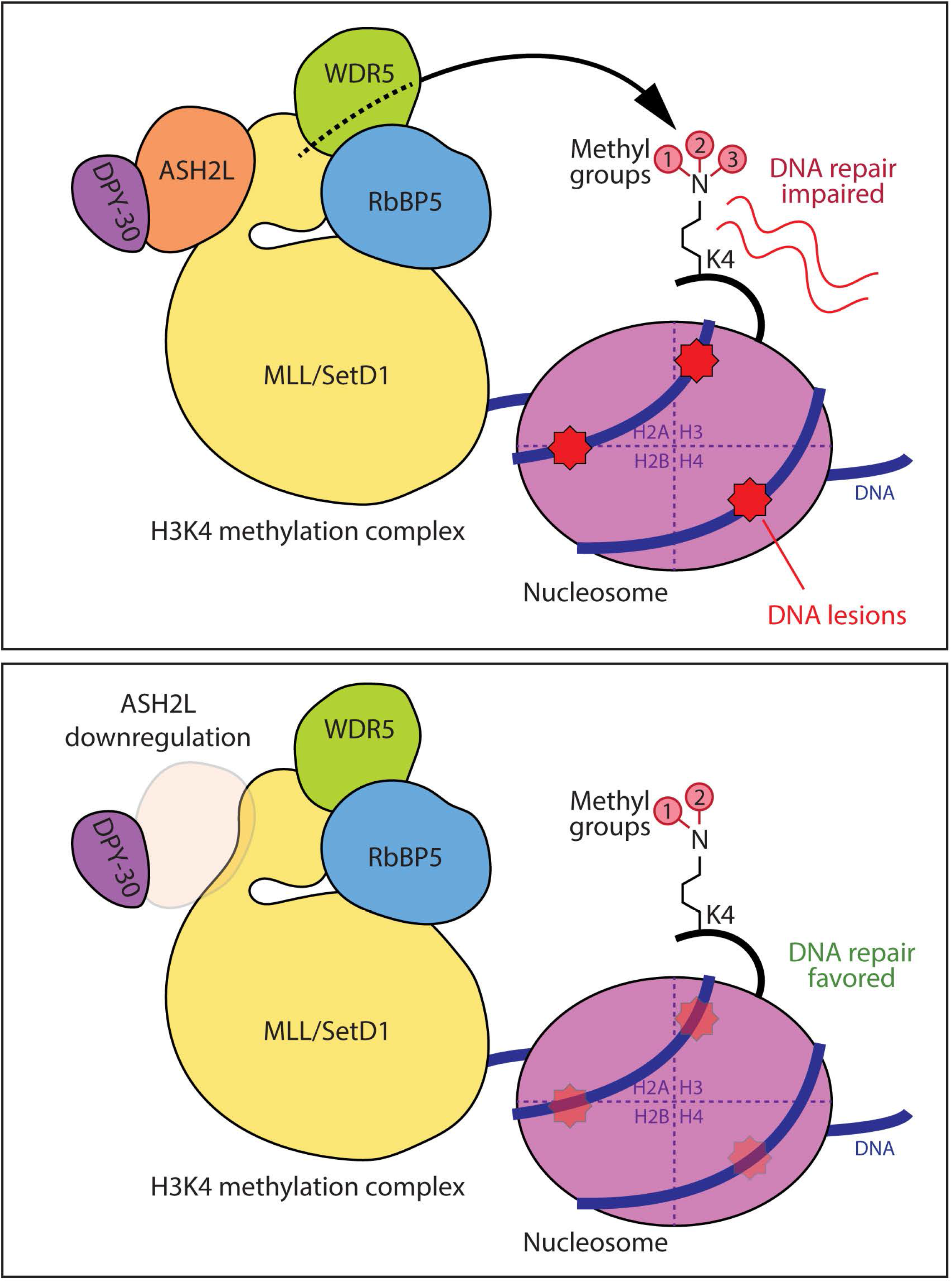
Schematic representation of the impact of ASH2L decrease on DNA repair. ASH2L is a core component of the H3K4 methylation complex involved in the addition of the third methyl on lysine 4 of histone 3 (H3K4). The trimethylation of H3K4 reduces the cell’s capacity to repair its DNA upon genotoxin-induced damage (28). In contrast, ASH2L silencing and the corresponding decrease in H3K4 trimethylation favors the ability of the cells to repair their damaged DNA (this study).

**Figure S6.**
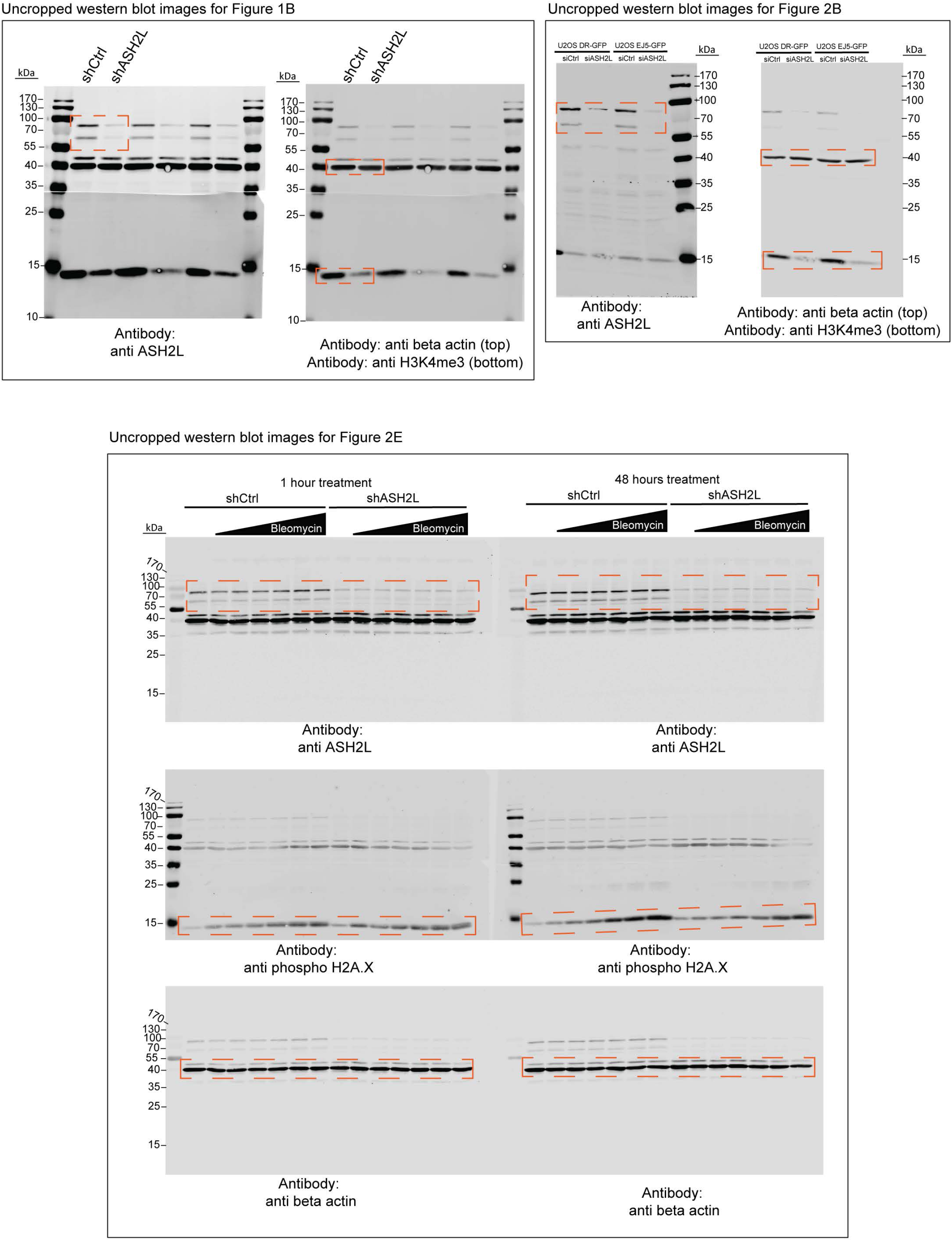
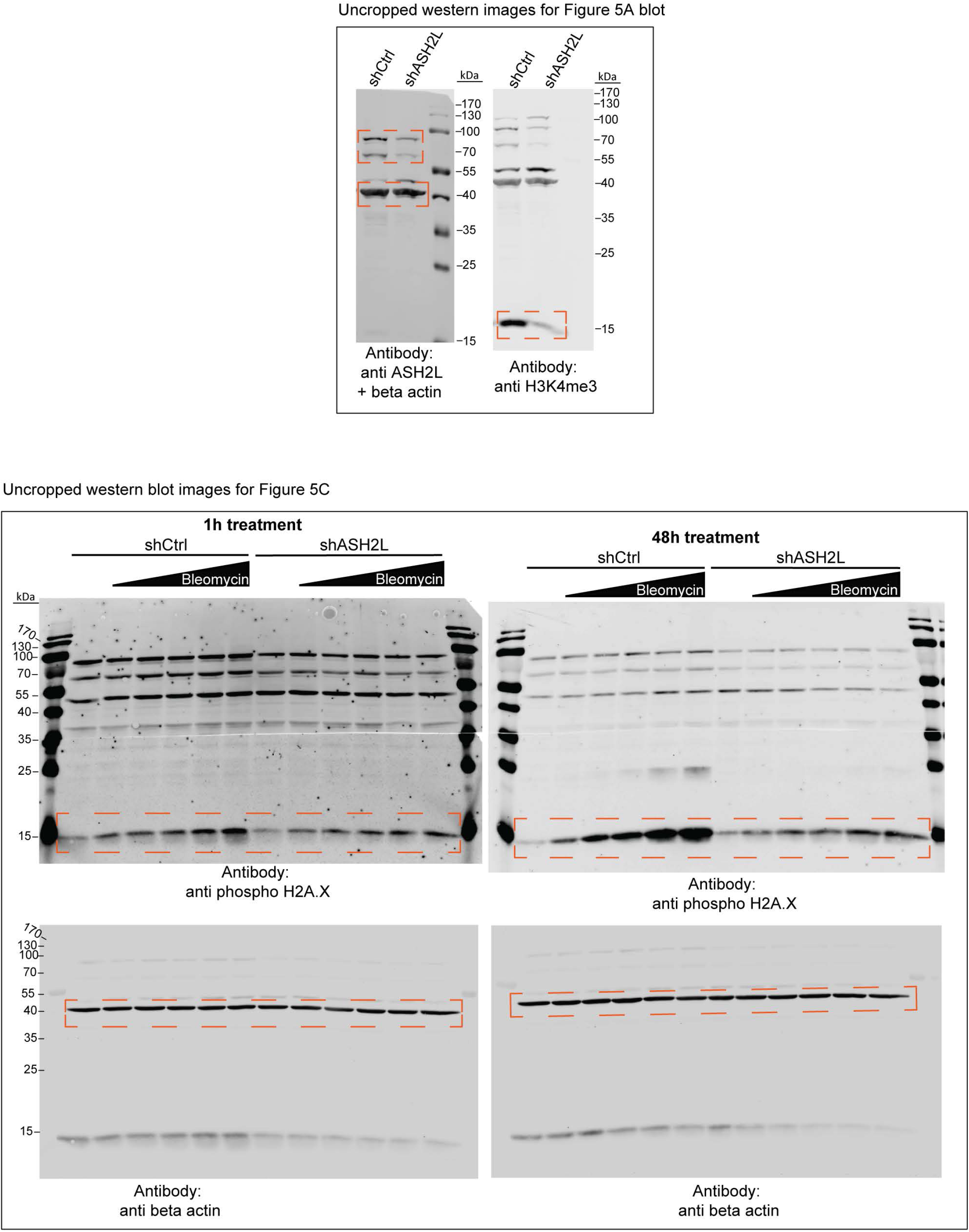
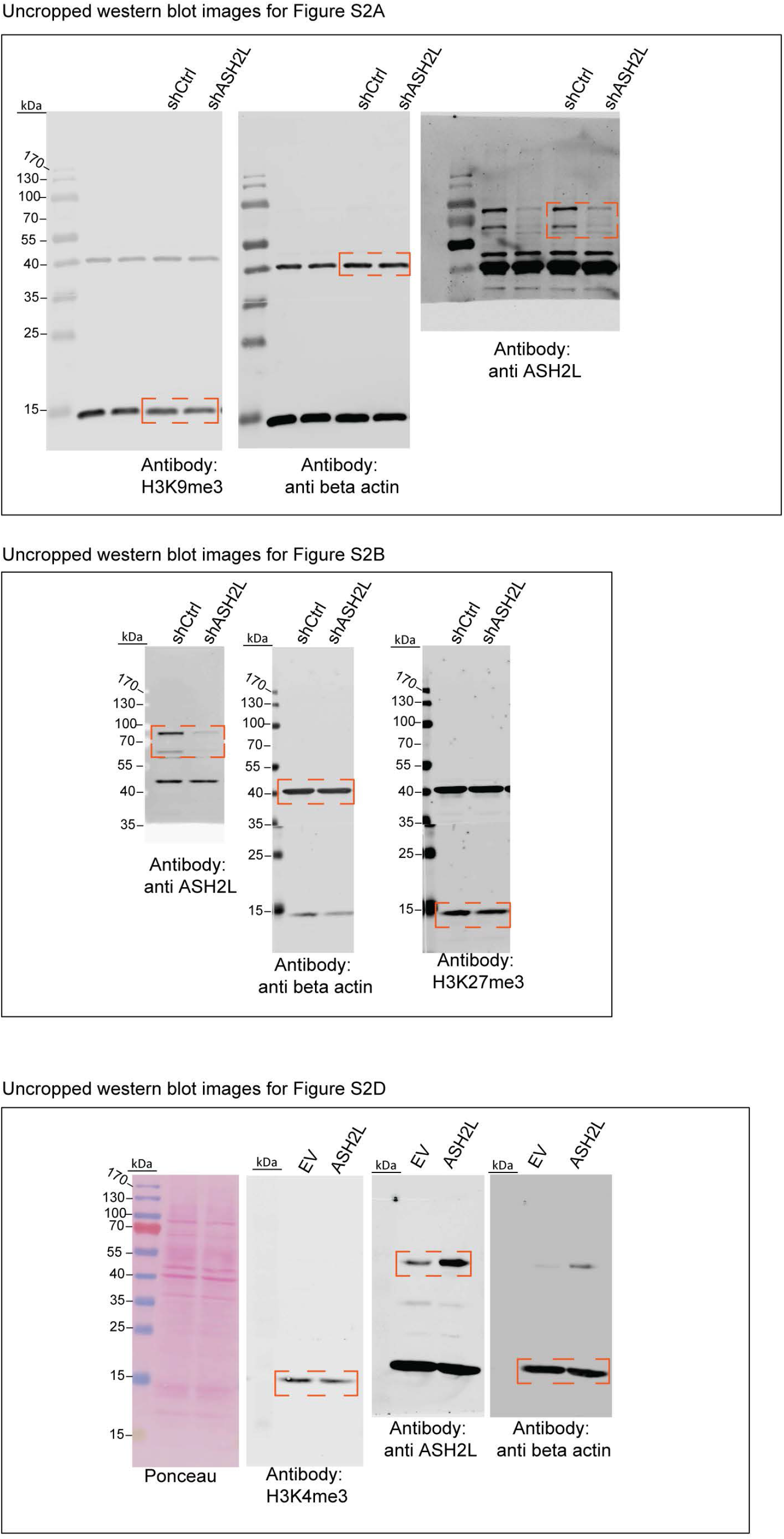
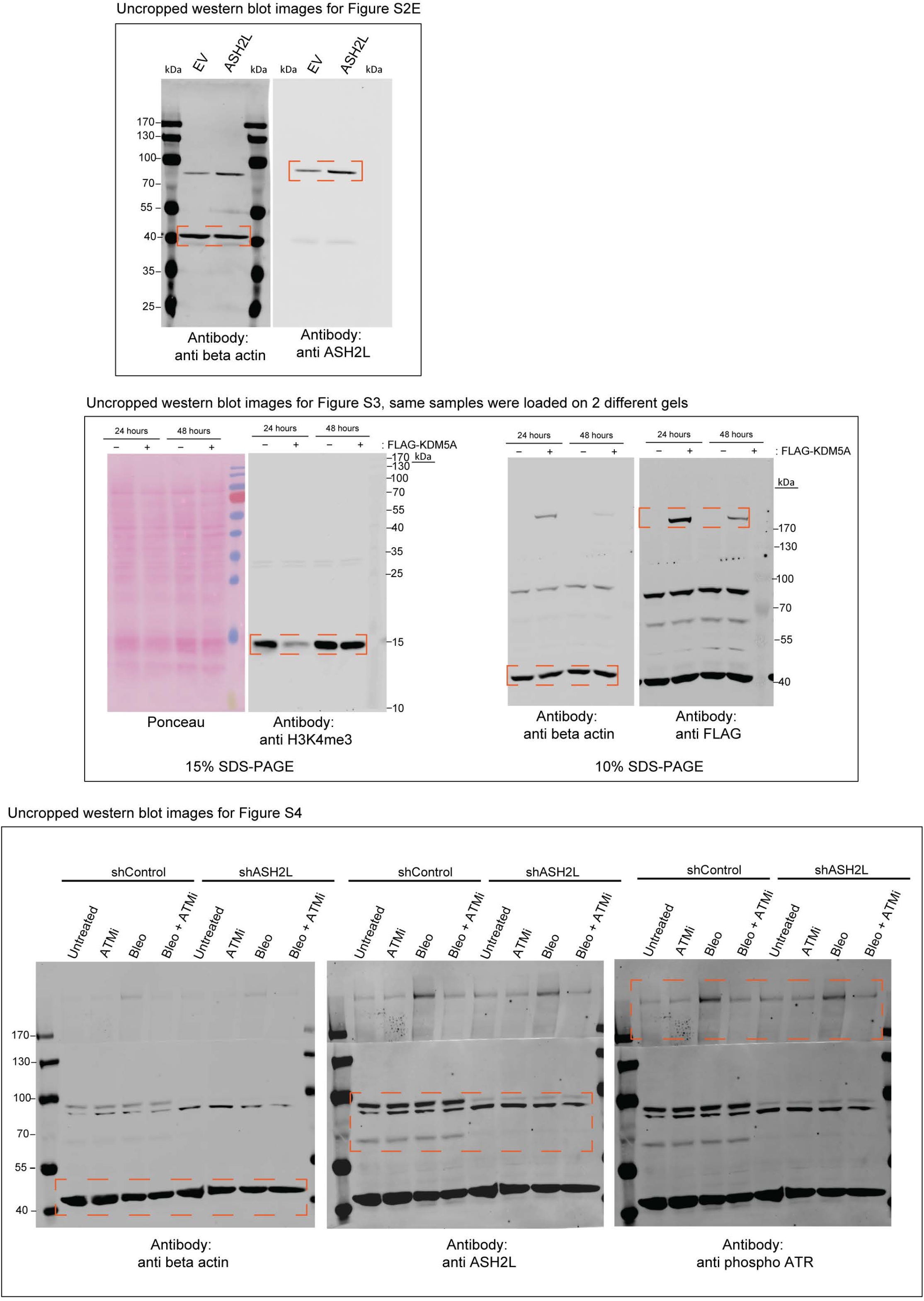
Uncropped western blots. The figure (in 4 parts) shows the uncropped blots that were used to generate the figures displaying western blot data.

